# Ameliorative potential of curcumin on cigarette smoke extract induced cognitive impairment in zebrafish

**DOI:** 10.1101/290536

**Authors:** Loganathan Thilagavathi, Sara Jabeen, Shreyas Belagod Ravishankar, Syed Shakeeb Ahmed, Thomas George, Narahari Rishitha, Arunachalam Muthuraman, Nallupillai Paramakrishnan

**Affiliations:** Department of Pharmacology, JSS College of Pharmacy, JSS Academy of Higher Education and Research, Mysuru-570 015, Karnataka, India

**Keywords:** Cigarette smoke extract, Cognitive disorder, Curcumin, *Danio rerio*, Montelukast

## Abstract

Curcumin is a major phyto-constituent of *Curcuma longa*. It has been reported that it that it showed various pharmacological actions via anti-oxidant, anti-inflammatory, and immunomodulatory actions. The present study undergoes the therapeutic evaluation of curcumin in cigarette smoke extract (CSE) exposure induced cognitive impairment in zebrafish. *Methods:* The cognitive impairment was induced by exposure to 25 ml of 200 % CSE; 10 minutes/day, for 7 consecutive days. The pre-treatment of curcumin (10 and 20 mg/kg) and montelukast(20 mg/kg) were exposed in 25 ml drug solution (10 minutes/day for 10 consecutive days). The changes of CSE induced cognitive dysfunction were evaluated by using different test methods such as colour recognition test; partition preference test; horizontal compartment test; and T-Maze tests. Further, the CSE exposure induced changes of biochemical parameters *i.e.,* acetylcholinesterase activity; lipid peroxidation; reduced glutathione; and total protein levels were estimated in the brain of zebrafish. The pre-exposure of curcumin shown to produce the ameliorative effect against CSE induced neurocognitive impairments along with alteration of biochemical changes. Theseresults were comparable to reference control *i.e.,* montelukast pre-treated group. Therefore, the curcumin can be served as newer medicines for immunological reaction associated neurocognitive disorders like Alzheimer and multiple sclerosis due to its potential anti-oxidative; anti-inflammatory; immunomodulatory; and acetylcholinesterase inhibitory actions.

**SUMMARY:** Loss of memory is a major problem in old age population. Curcumin used to treats the various neurological disorders. Curcumin possess the ameliorative potential in toxin induced neurocognitive function.

## INTRODUCTION

Curcumin is one of the primary active constituents in *Curcuma longa*. In India, rhizome part of this plant widely used for the various food preparations as nutrients and coloring agents. It is a herbaceous perennial plant, and it belongs to *Zingiberaceae* family (Sumathi et al., 2017). Traditionally Ayuervedic, Siddha and Unani systems of medicine are recognized as turmeric can be a cure in chronic illness disorders; and it is prophylaxis for multiple infections, cancer cell growth; cardioprotection as well as neuroprotection (Elufioye et al., 2017; Sen and Chakraborty, 2017). Currently, the active phytoconstituent *i.e.,* curcumin is documented that, it possesses the potential neuroprotective action via multiple pharmacological mechanisms like anti-oxidant; anti-inflammatory (Sharma and Nehru, 2018; Zhang and Li, 2017); pro-apoptotic action on cancer cell (Rivera et al., 2017); anti-apoptosis in endothelial cells (Loganes et al., 2017); direct regulation of cytokines (Oh et al., 2018); reduce the mast cell activation and release of histamine (Li et al., 2014); activation of potassium ion channel (Chen et al., 2015)and inhibition of Transient Receptor Potential Melastatin 2 (TRPM2) channel (Kheradpezhouh et al., 2016); cholinergic modulatory action (Akinyemi et al., 2017); and inhibition of acetylcholinesterase activity (Kalaycioglu et al., 2017; Tello-Franco et al., 2013). In addition, it is also reported that it reduces the accumulation of β-amyloid peptide via reduction of β-amyloid synthesis and enhance the β-amyloid catabolic process in the brain and spinal cord regions(Fan et al., 2017; Zheng et al., 2017). Due to its pleiotropic function on cellular and molecular mechanism of central nervous system (Karuppagounder et al., 2017; Luthra and Lal, 2016); it is expected to produce the neuroprotective action against multiple pathophysiological events such as hyperglycemia (Daugherty et al., 2018); hypoxia (Shen and Yu, 2008); trauma (Qi et al., 2017); and toxin like picrotoxin, streptozotocin and pentylenetetrazole (Bassani et al., 2017; Choudhary et al., 2013; Wang et al., 2017); and cigarette smoke(Saito et al., 2018).

Now a day, the exposure to cigarette smoke in the environment affects the multiple neuroendocrinological functions (Jamal et al., 2018; Saito et al., 2018); and it alters the genetic materials leads to affects the neurobehavioural function (Dogan et al., 2017). In addition, the prevalence rate of cigarette smoking (CS) induced cognitive impairments is higher in developing countries (Ashare et al., 2014; Bashir et al., 2017). There is no specific medicaments are available for the management and of CSE associated neurotoxicity (Beckham et al., 2018). Day by day, CS induced cognitive changes and other neurological problem raises the great challenges in the field of medical research(Giordano et al., 2018). CS is posses the more than 60 cytotoxic chemicals. Some of the chemicals (*i.e.,* carbon monoxide; nitrosamines; aryl hydrocarbon; phenolic compounds; and polynuclear aromatic compounds) are potent modulators of neuroendocrine function; and inducer of neurodegeneration & neuronal death (Ichitsubo and Kotaki, 2018; Tweed et al., 2012; Yun et al., 2018). In addition, CS is attacking the nervous system via free radical generation; activation of inflammatory cytokines and chemokine functions; enhances the immunological reactions via mast cell activation; and dysfunction of mitochondria; endoplasmic reticulum and nucleus of the cells; which in turn cause the oxidative stress; apoptosis; neurodegeneration and neuronal death(Chan et al., 2016). Therefore, newer research is needed to explore the better medicaments for the protection of the neurological system from neurotoxic agents and prevention of CS induced neurocognitive disorders. Zebrafishare widely used for the assessment of neurocognitive behaviours (Massarsky et al., 2018; Sharma et al., 2017). Montelukast is a potent anti-inflammatory and immunosuppressive drugs, andit is used for the treatment of bronchial asthma. It protects the neurological system via blocking of cysteinyl-leukotriene receptor and it also produces the improvements of cognitive functions against various neurotoxic events (Grinde and Engdahl, 2017; Zhang et al., 2016). And it was selected for a reference control group in this study. Thus, the present study, designed to investigate the therapeutic potential of curcumin on cigarette smoke extract (CSE) induced cognitive impairment in zebrafish.

## MATERIALS AND METHODS

### Animal

Wild type adult (< 8 month old male) zebrafish was used in this study. The animal kept in 10 liters (potable water) housing tank. The tank was maintained with aerator for aeration process; the temperature, *i.e.,* 25 ± 2°C was maintained and 11 to 12 hours light and dark cycle of photoperiod to maintain the normal circadian rhythmicity. All the animals were acclimatized for 2 weeks before the study. All behavioral observation were performed between the 09.00 AM to 01.00 PM to avoid the hormone associated neurobiological interaction and neurobehavioral abnormalities.

### Preparation cigarette smoke extract (CSE)

The 200 % CSE was prepared by the described method of Yoon et al. (2013) with minor modification. Briefly, the commercial cigarettes (Gold Flake packet purchased from Mysuru, India) were used in this study; and each cigarette contained 12□mg tar and 1.2□mg nicotine. Six pieces of cigarettes were placed in the smoke-pump machine and smoked continuously. The cigarette smoke was allowed to saturate in 100□ml of pre-warmed distilled water. This solution was passed through a 0.22 μm pore filter (Millipore Corporation, Maharashtra, India) to remove large particulates and bacteria. The solution was considered as 200 % CSE and used for the further experiments for the neurocognitive disorder.

#### Induction of cognitive impairment by CSE exposure

The cognitive impairment was induced by exposing of cigarette smoke extract (CSE) in zebrafish as a descriptive method of Folkesson et al. (2016); and Massarsky et al. (2018). Our pilot study revealed that the mortality started at 30 minutes exposure with 200 % of CSE. This experiment, each zebrafish was exposed in 50 ml beaker containing 25 ml of 200 % CSE for 10 minutes/day, for 7 consecutive days.

### Experimental protocol

In this study, the experimental design was made with five groups of adult male zebrafish (n = 20). Group I was served as a normal control group. The group II was employed as CSE (200 %; 10 minute/day, for 7 consecutive days) challenged group. Group III and IV were served as pre-exposure of curcumin (10 and 20 mg/kg; 10 minute/day, for 7 consecutive days) against CSE challenge. The curcumin exposures were made 20 minutes before the CSE challenge in zebrafish animals. The group V was served as pre-exposure of montelukast (20 mg/kg; 10 minute/day, for 7 consecutive days) against CSE challenge. The montelukast was served as a reference control group. The behavioral parameters were assessed on the 8^th^ day; whereas, T maze training was performed on the 8^th^ day and assessment of memory function performed on day 9. On the 9^th^ day, all the animals were sacrificed and brain samples were collected for the biochemical estimation. Experimental design of this study illustrated in figure 1.

**Figure 1.**
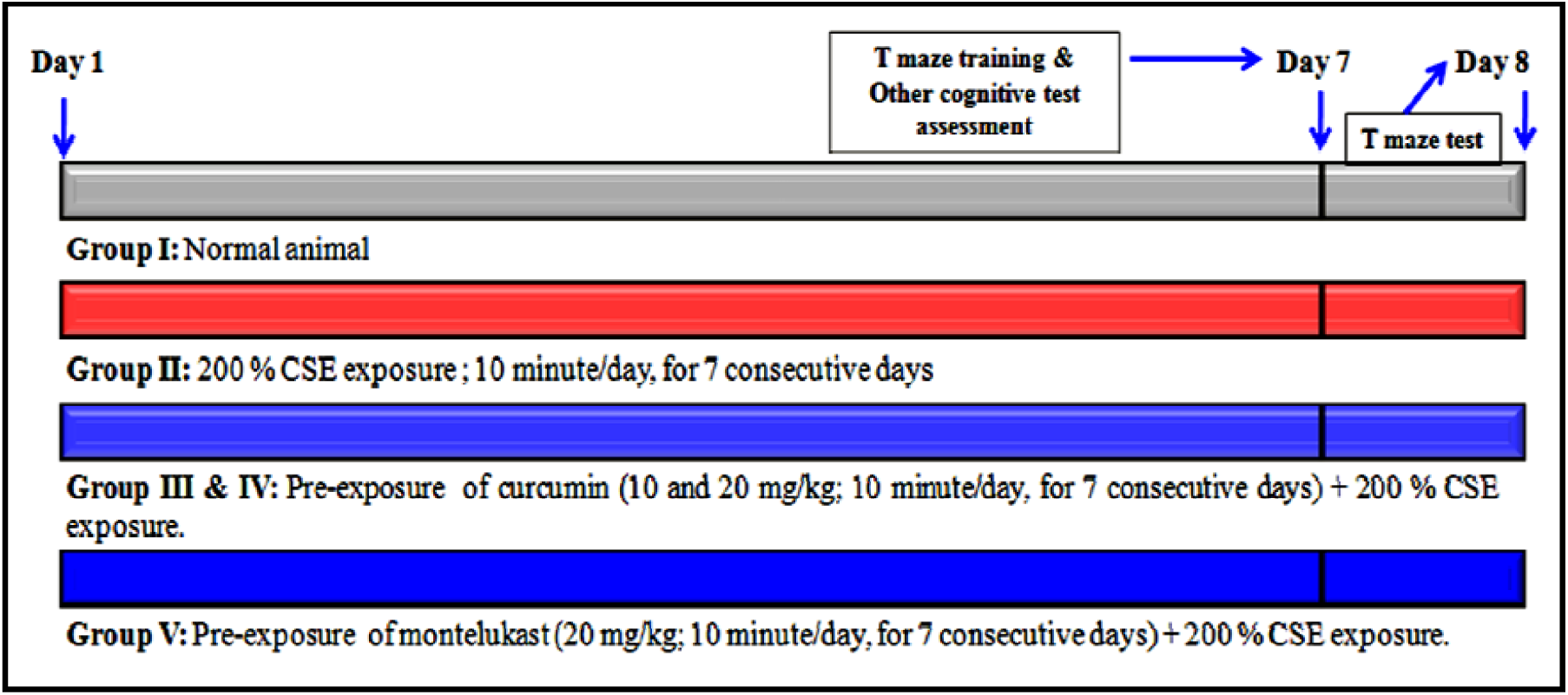
Experimental design for the evaluation of curcumin in CSE induced changes of cognition.

### Assessment of cognitive function

The cognitive functions are evaluated with different tests like Color recognition test; partition preference test; horizontal compartment tests; and T-Maze tests. Zebrafish was engaged in the evaluation of neurocognitive functions with different test apparatus after exposing of CSE exposure. All animals were acclimatized for 5 minutes before in all behavioral test apparatus to minimize the error report. The animal movements were tracked by USB camera with computer software (12 Megapixel USB Camera, Intex, India).

#### Color recognition test

Color recognition test is one of the methods of spatial memory function as a descriptive method of Dubey *et al.* (2015) with minor modification of Rishitha and Muthuraman (2018). Briefly, the test chamber (20L × 10W × 10H, cm; Length, Width, and Height respectively) was divided into two equal parts. One half of the chamber (10L × 10W × 10H, cm) is covered with red color glass and it is considered as the punishment chamber. Another half of the chamber is covered with green colored glass, and it is considered as the reward chamber. The water level was maintained up to the 5 cm. The cognitive function was evaluated by assessing *“Time spent in the green chamber”* termed as TSGC and *“Number of entries to the red chamber”* termed as NERC. When animal placed in the center of the chamber it prefers the green side; it indicates the presence or improvement of memory function. If the animaldoes not prefer green chamber it has a poor memory.

#### Partition preference test

The Partition preference test is another method for the assessment of working and spatial memory function as a descriptive method of Dubey *et al.* (2015) with minor modification of Rishitha and Muthuraman (2018). Briefly, the chamber (20L × 10W × 10H; cm) water level was maintained up to the 3 cm; it is divided into two equal (10L × 10W × 10H; cm) parts with vertical glass slide movement plate and the gap was maintained for 1 cm from the bottom of the inner glass chamber. Right side chamber considered as home chamber; whereas, left side chamber consider as target chamber. The cognitive function was evaluated by assessment of a number of the entry to the target chamber and calculated as *“Percentage entry to the target chamber”* (% ETC), and another one is named as *“Time spent in the target chamber”* (TSTC). When animal placed in the home chamber; it prefers to reach the target chamber. If the animaldoes not enter into a home chamber within 60 seconds time period; the animal was guided to reach the target chamber and kept in same target chamber for a further 20 seconds by the closing of entry site with a glass slide. The preference of both chambers indicates the enhanced memory function. The preference of home chamber is the lack of memory function.

#### Horizontal compartment test

The Horizontal compartment test is one of the methods for the assessment of neurocognitive function and it is described by Dubey *et al.* (2015) with minor modification of Rishitha and Muthuraman (2018). Briefly, the chamber (20L × 10W × 24H; cm) water level was maintained up to the 21 cm; it is divided into three horizontal (7 × 7 × 7; cm height) compartments by marking a line in the outer chamber. One day before, separate training was given (120 seconds) to all fishes to swim in the all compartments; if the animaldoes not preferall compartments, animals were guided with guiding tool and food pellets. Next day, the cognitive functions were evaluated by assessment of “*time spent in the upper segment”* (TSUS); and “*time spent in the lower segment”* (TSLS). Generaly, when animal placed in the test chamber; it prefers to swim in the upper segment of the chamber within 15 seconds. It indicates that the animal has a normal or improved memory function. If the animal does not prefer the upper segment; and animals were swim in middle or lower segment indicates that loss of memory.

#### T-Maze tests

The t-maze test is one of the established methods for the assessment of rodent neurocognitive function. The various researchers modified the T-maze test apparatus, and it is shown reproducible result in zebrafish. T-Maze tests for zebrafish described by Colwill *et al.* (2005) with minor modification of Muthuraman and Rishitha (2018). Briefly, the T maze apparatus consists of two short arms (10Lx 6W × 10H; cm) with a different color (one arm with red color glass; and another end arm with green color glass). And one long arm (20L × 10W × 10H; cm) covered with home chamber (5L × 6W × 10H; cm); and it is made with normal non-transparent glass. The green color arm employed as a favourable environment with the reward of a food pellet, and another red color arm employed as punishment by a string with a glass rod. One day before the test assessment; all animals were allowed to learn the T maze environment; and allow entering in the green chamber. If not learned, animals were guided to reach the green chamber. Next day, animals were placed in the corner of long arm *i.e.,* starting point from home chamber and target point was identified as an entry to any one of the short arm. The starting point and target point were separated by vertical slide control panel. Each fish was explored for 2 minutes for the learning and memory assessment. The transfer latency (TL) and percentage target (green) chamber preference (% TCP) were noted for the assessment of cognitive function.

### Estimations of biomarker changes

After assessment of behavioral assessment, the zebrafish brain samples were isolated immediately by the microsurgical method and freeze dried at -4 °C. Next day, all samples were homogenated with phosphate buffer solution. The supernatant was collected by centrifugation at 1372 g-force for 15 minutes. The supernatant of zebrafish brain samples was used for estimation of tissue biomarker changes, *i.e.,* acetylcholinesterase (AChE) activity; lipid peroxidation (LPO); reduced glutathione (GSH); and total protein levels.

#### Estimation of AChE level in the zebrafish brain

The AChE activity level of zebrafish brain samples was estimated by a spectroscopic method as described by Ellman *et al.* (1961) with minor modification of Rishitha and Muthuraman (2018). Briefly, 500 µl of zebrafish brain supernatant was mixed with 0.25 ml of DTNB (0.001 M) and incubated for 10 minutes. The formation of yellow color chromogen products and their colour intensity were assessed with the observation of absorbance changes by using a spectrophotometer (DU 640B Spectrophotometer, Beckman Coulter Inc., CA, USA) at 420 nm wavelength. This absorbance changes varied corresponding to the changes acetylthiocholine hydrolyzed product by AChE activity. Further, the AChE activity levels were calculated by using the following formula *i.e.,* R = δ O.D X Volume of the assay (3 ml) / ε X mg of protein. Hence, R is representing the rate of enzyme activity in ‘n’ mole of acetylthiocholine iodide hydrolyzed per minute per mg protein. δ O.D. is representing a change of absorbance per minute. ε is representing the extinction coefficient value *i.e.,* 13600 per mole per centimeter. The results were expressed as μM of acetylthiocholine hydrolyzed per milligram of protein per minute.

#### Estimation of lipid peroxidation product level in the zebrafish brain

The lipid peroxidation (LPO) product level of zebrafish brain samples was estimated by a spectroscopic method as described by Ohkawa *et al.* (1979) with minor modification of Rishitha and Muthuraman (2018). Briefly, 0.2 ml of zebrafish brain tissue supernatant was mixed with 0.2 ml of 8.1% w/v of sodium dodecyl sulphate (SDS), 1.5 ml of 30 % v/v of acetic acid (pH 3.5), 1.5 ml of 0.8 % w/v of thiobarbituric acid and volume made upto 4 ml of distilled water. The reaction mixtures were incubated at 95 °C for 1 hour. Then rapidly cooled with tap water and further 1 ml of distilled water and 5 ml of n-butanol-pyridine (15:1 v/v) mixture were added. After 10 minutes, tubes were centrifuged at 1372 g-force for 15 min. The formation of pink color chromogen products and their colour intensity were assessed with the observation of absorbance changes by using spectrophotometer(DU 640B Spectrophotometer, Beckman Coulter Inc., CA, USA) at 535 nm wavelength. These absorbance readings were used for the further calculation of LPO levels. The results were expressed as nM per mg of protein.

#### Estimation of GSH level in the zebrafish brain

The GSH level of zebrafish brain samples was estimated by a spectroscopic method as described by Ellman (1959) with minor modification of Rishitha and Muthuraman (2018). Briefly, 0.5 ml supernatant was mixed with 2 ml of disodium hydrogen phosphate solution (0.3 M) and 0.25 ml of freshly prepared DTNB solution (0.001 M). The formation of yellow color chromogen products and their colour intensity were assessed with the observation of absorbance changes by using a spectrophotometer (DU 640B Spectrophotometer, Beckman Coulter Inc., CA, USA)at 412 nm wavelength. These readings are used for the further calculation of GSH levels in tissue samples. The results were expressed as μM of GSH / mg of protein.

#### Estimation of total protein level in the zebrafish brain

The total protein level of zebrafish brain samples was estimated by a spectroscopic method as described by Lowry *et al.* (1951) with minor modification of Rishitha and Muthuraman (2018). Briefly, 300 µl of zebrafish brain supernatants were diluted with distilled water upto 1 ml. Further, 5 ml of Lowry’s reagent was mixed with the solution and allow standing for further 15 minutes at room temperature (37 °C). Then 0.5 ml of Folin-Ciocalteu reagent was added slowly and vortexed vigorously at room temperature for 30 min. The formation of purple color chromogen products and their color intensity were assessed with the observation of absorbance changes by using spectrophotometer (DU 640B, UV-Spectrophotometer, Beckman Coulter Inc., CA, USA) at 750 nm wavelength. These readings were used for the further calculation of total protein levels in tissue samples. The results were expressed as mg of protein per ml of supernatant.

### Statistical analysis

All the results were expressed as the mean ± standard deviation (SD). Data obtained from all behaviour tests and tissue biomarkers were statistically analyzed using one-way analysis of variance (ANOVA) followed by Tukey’s test were applied for Post-hoc analysis by using Graph pad prism Version-5.0 software. A probability value of *p* < 0.05 was considered to be statistically significant.

## RESULTS

### Effect of curcumin in CSE induced changes of colour recognition test

The exposure of CSE (200 %) produce the significant (*p* < 0.05) cognitive impairment as an indication of decreasing TSGC and raising NERC values when compared to a normal control group. The pre-exposure of curcumin (10 and 20 mg/kg) shown to produce the neuroprotective action against CSE exposure induced cognitive impairment in a dose-dependent manner. Furthermore, the pre-exposure of reference compound, *i.e.,* montelukast (20 mg/kg) produced the significant ameliorative action against CSE induced cognitive impairment changes. It indicates that the pre-exposure of curcumin produce the neuroprotective action against CSE toxicity. The results are illustrated in figure 2a and 2b.

**Figure 2.**
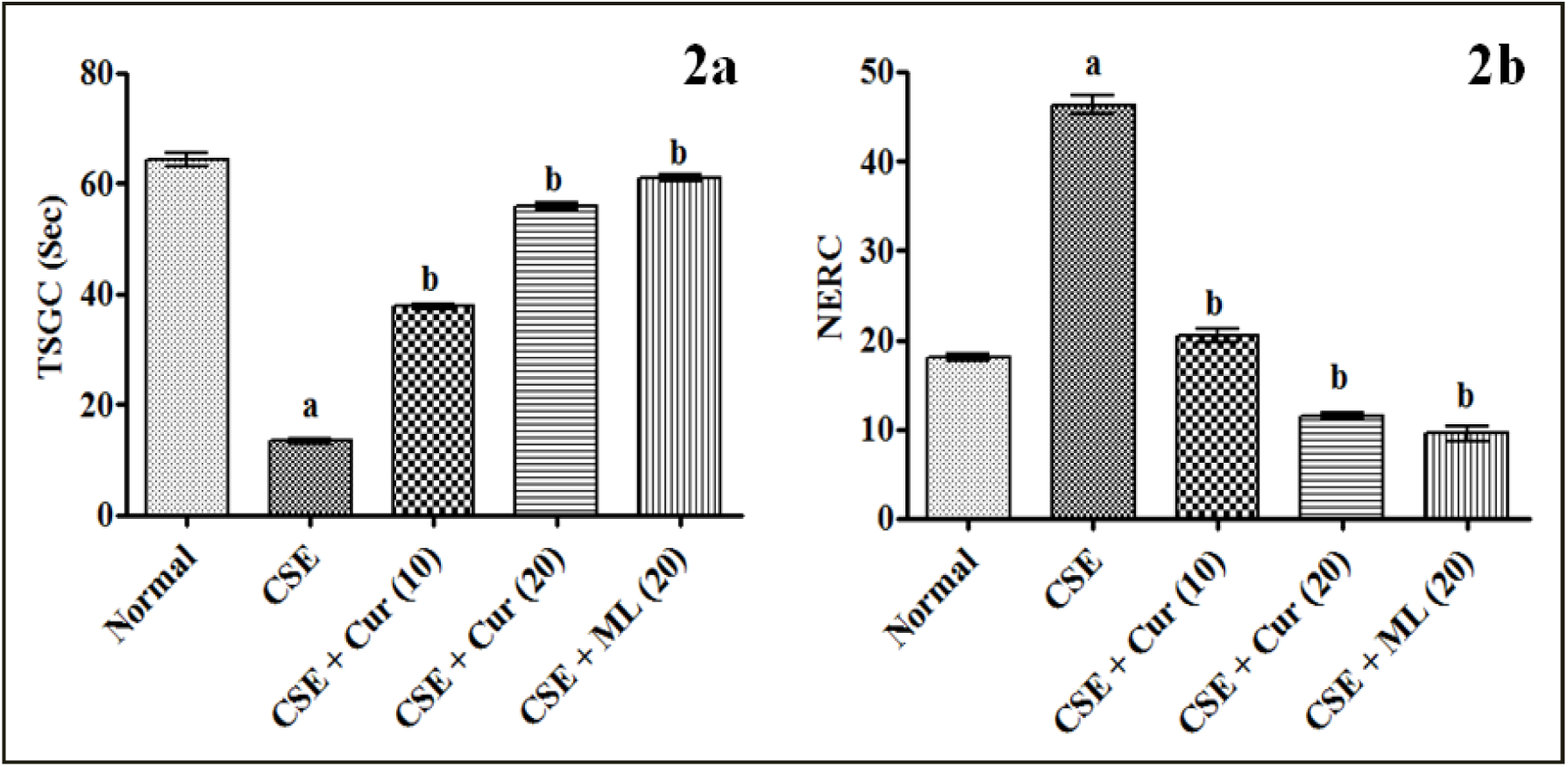
Effect of curcumin in CSE induced changes of color recognition test. Digits in parenthesis indicate dose mg/kg. Data were expressed as mean ± SD, n = 6 zebrafish per group. ^a^*p* < 0.05 Vs normal group. ^b^*p* < 0.05 Vs CSE exposure group. Abbreviation: CSE, cigarette smoke extract; Cur, curcumin; ML, montelukast; and TSGS, time spent in green chamber; NERC, number of entry to the red chamber.

### Effect of curcumin in CSE induced changes of partition preference test

The exposure of CSE (200 %) produce the significant (*p* < 0.05) cognitive impairment as an indication of decreasing % ETC and TSTC values when compared to a normal control group. The pre-exposure of curcumin (10 and 20 mg/kg) shown to produce the neuroprotective action against CSE exposure induced cognitive impairment in a dose-dependent manner. Furthermore, the pre-exposure of reference compound, *i.e.,* montelukast (20 mg/kg) produced the significant ameliorative action against CSE induced cognitive impairment changes. It indicates that the pre-exposure of curcumin produce the neuroprotective action against CSE toxicity. The results are illustrated in figure 3a and 3b.

**Figure 3.**
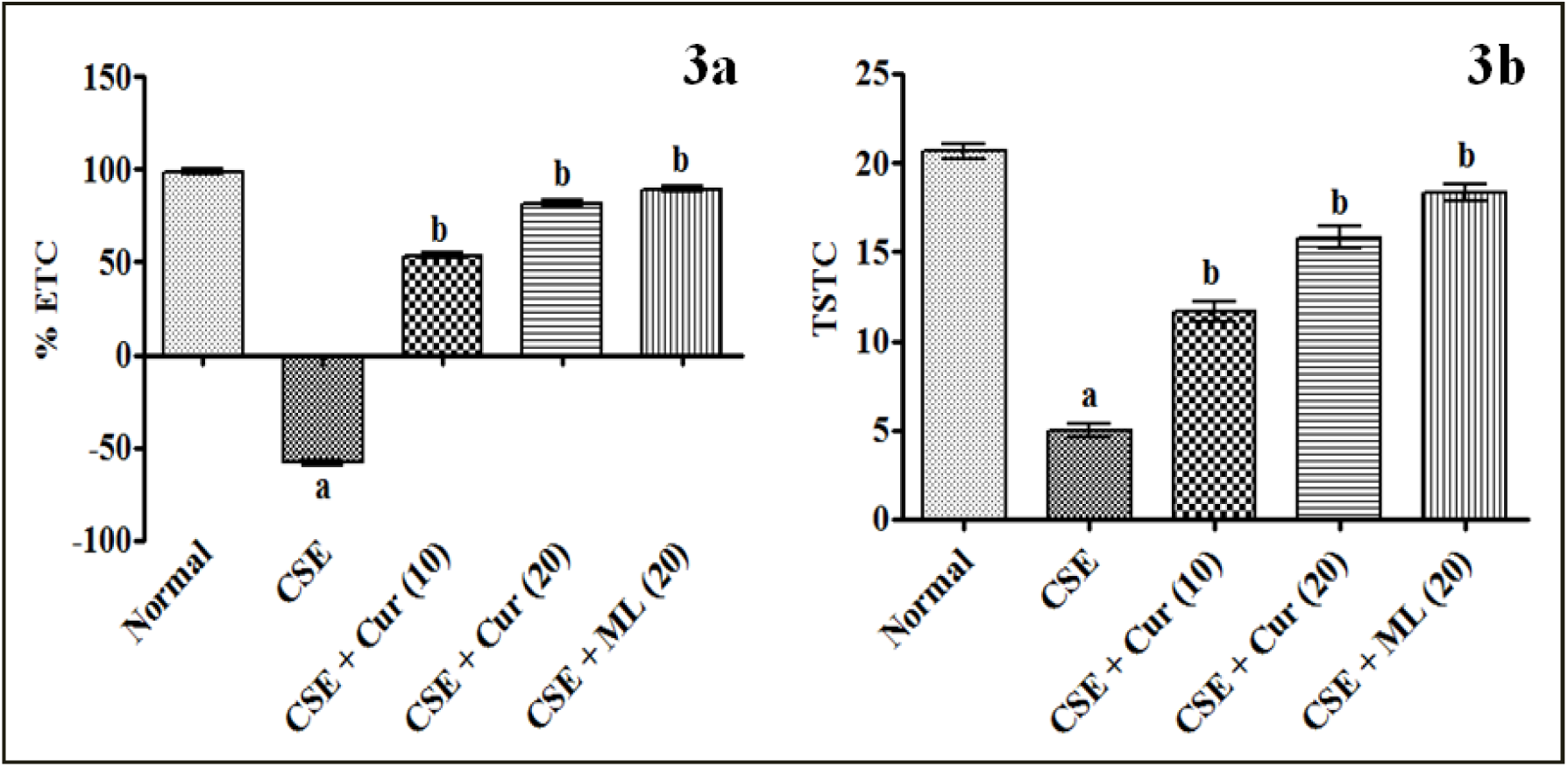
Effect of curcumin in CSE induced changes of partition preference test. Digits in parenthesis indicate dose mg/kg. Data were expressed as mean ± SD, n = 6 zebrafish per group. ^a^*p* < 0.05 Vs normal group. ^b^*p* < 0.05 Vs CSE exposure group. Abbreviation: CSE, cigarette smoke extract; Cur, curcumin; ML, montelukast; % ETC, percentage entry to the target chamber; and TSTC, time spent in the target chamber.

### Effect of curcumin in CSE induced changes of horizontal compartment test

The exposure of CSE (200 %) produce the significant (*p* < 0.05) cognitive impairment as an indication of decreasing TSUS and increasing the TSLS values when compared to a normal control group. The pre-exposure of curcumin (10 and 20 mg/kg) shown to produce the neuroprotective action against CSE exposure induced cognitive impairment in a dose-dependent manner. Furthermore, the pre-exposure of reference compound, *i.e.,* montelukast (20 mg/kg) produced the significant ameliorative action against CSE induced cognitive impairment changes. It indicates that the pre-exposure of curcumin produce the neuroprotective action against CSE toxicity. The results are illustrated in figure 4a and 4b.

**Figure 4.**
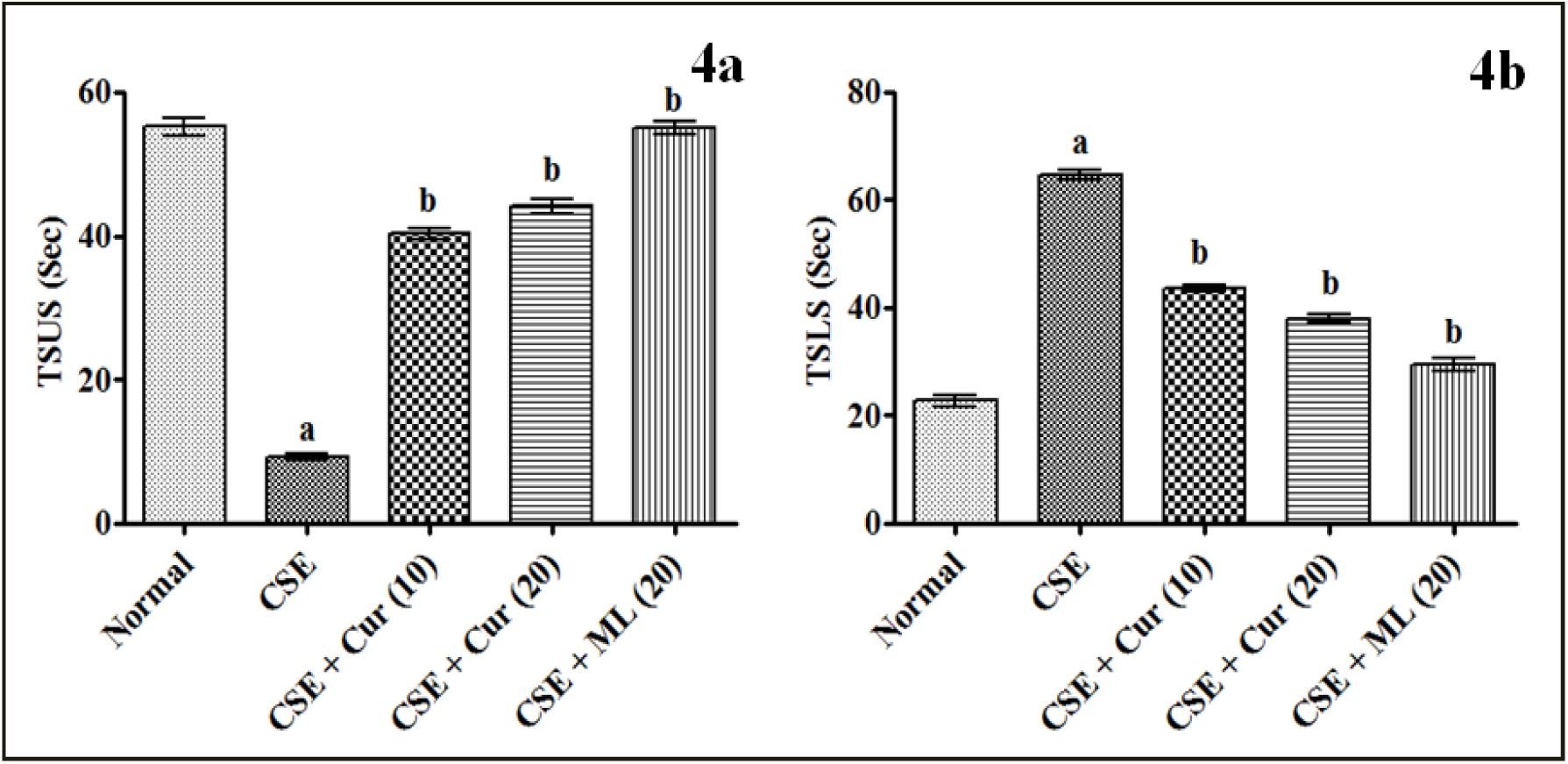
Effect of curcumin in CSE induced changes of horizontal compartment test. Digits in parenthesis indicate dose mg/kg. Data were expressed as mean ± SD, n = 6 zebrafish per group. ^a^*p* < 0.05 Vs normal group. ^b^*p* < 0.05 Vs CSE exposure group. Abbreviation: CSE, cigarette smoke extract; Cur, curcumin; ML, montelukast; TSUS, time spent in the upper segment; and TSLS, time spent in the lower segment.

### Effect of curcumin in CSE induced changes of T-Maze tests

The exposure of CSE (200 %) produce the significant (*p* < 0.05) cognitive impairment as an indication of increasing TL and decreasing % TCP values when compared to a normal control group. The pre-exposure of curcumin (10 and 20 mg/kg) shown to produce the neuroprotective action against CSE exposure induced cognitive impairment in a dose-dependent manner. Furthermore, the pre-exposure of reference compound, *i.e.,* montelukast (20 mg/kg) produced the significant ameliorative action against CSE induced cognitive impairment changes. It indicates that the pre-exposure of curcumin produce the neuroprotective action against CSE toxicity. The results are illustrated in figure 5a and 5b.

**Figure 5.**
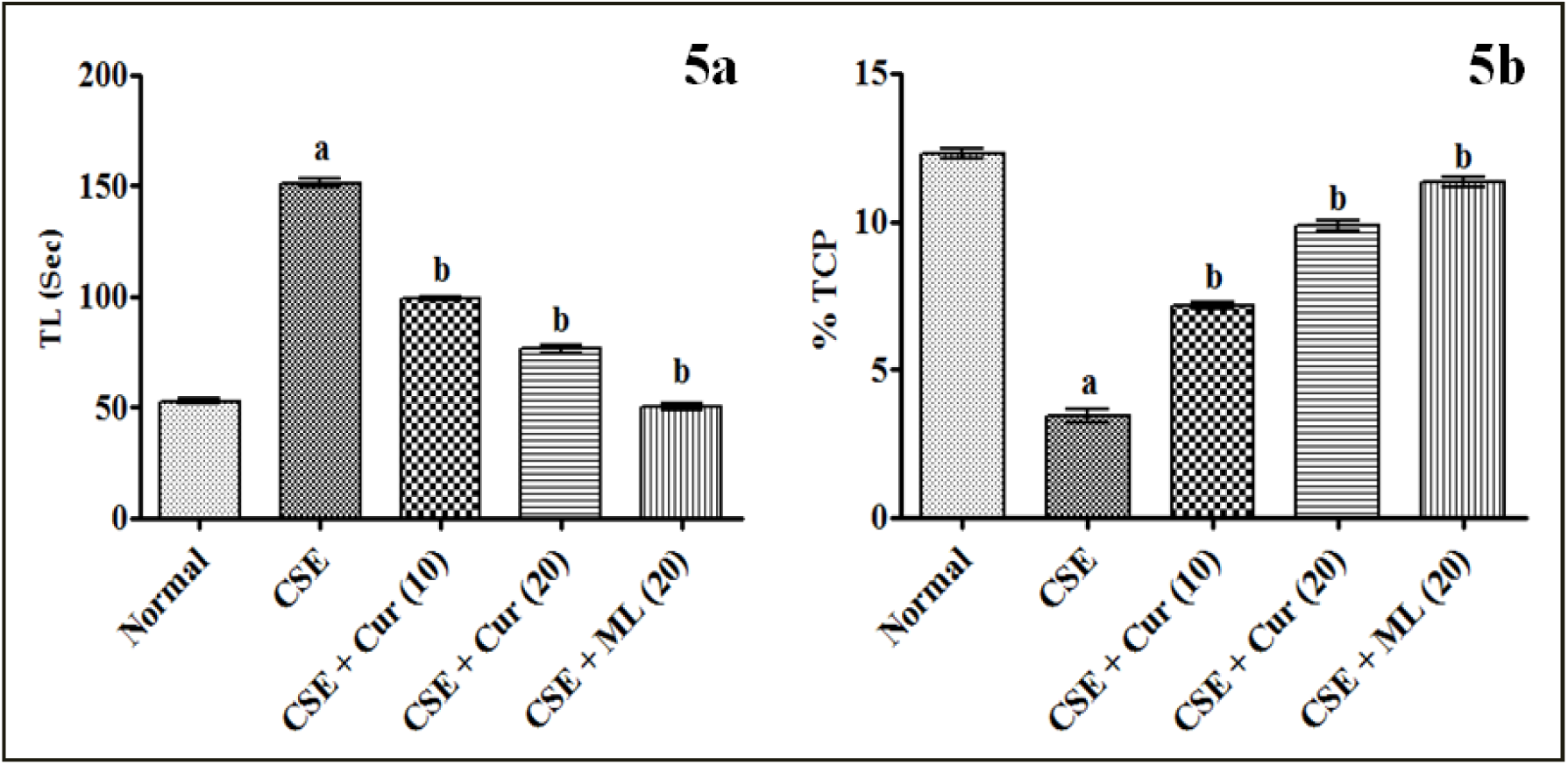
Effect of curcumin in CSE induced changes of T-Maze tests. Digits in parenthesis indicate dose mg/kg. Data were expressed as mean ± SD, n = 6 zebrafish per group. ^a^*p* < 0.05 Vs normal group. ^b^*p* < 0.05 Vs CSE exposure group. Abbreviation: CSE, cigarette smoke extract; Cur, curcumin; ML, montelukast; TL, transfer latency; and % TCP, percentage target (green) chamber preference.

### Effect of curcumin in CSE induced zebrafish brain biomarkers changes

The exposure of CSE (200 %) produce the significant (*p* < 0.05) rising of brain AChE activity and LPO, and decrease the GSH level when compared to a normal control group. The pre-exposure curcumin (10 and 20 mg/kg) shown to produce the attenuation effect against CSE exposure induced biochemical changes in a dose-dependent manner. Further, the pre-exposure of reference compound, *i.e.,* montelukast (20 mg/kg) produced the significant ameliorative effect against CSE induced tissue biomarker changes in zebrafish brain samples. It indicates that the pre-exposure of curcumin produce the neuroprotective action against CSE induced cognitive dysfunction via free radical scavenging; anti-lipidperoxidative; and reduction of acetylcholinesterase activity. The results are tabulated in table 1.

**Table 1.** Effect of curcumin in CSE induced zebrafish brain biomarkers changes.

| **Groups** | **AChE activity**<br>($\mu\text{M}$ / mg of protein / min) | **GSH**<br>($\mu\text{M}$ / mg of protein) | **LPO**<br>(nM / mg of protein) |
| --- | --- | --- | --- |
| Normal | $17.2 \pm 1.72$ | $16.6 \pm 2.17$ | $4.9 \pm 1.02$ |
| CSE | $51.3 \pm 1.05^a$ | $5.8 \pm 1.02^a$ | $13.8 \pm 1.17^a$ |
| CSE + Cur (10) | $32.9 \pm 0.98^b$ | $11.3 \pm 1.52^b$ | $10.2 \pm 0.69^b$ |
| CSE + Cur (20) | $23.4 \pm 1.16^b$ | $13.5 \pm 1.09^b$ | $6.8 \pm 0.72^b$ |
| CSE + ML (10) | $18.9 \pm 1.28^b$ | $15.6 \pm 1.39^b$ | $5.2 \pm 0.53^b$ |

Digits in parenthesis indicate dose mg/kg. Data were expressed as mean ± SD, n = 6 zebrafish per group. <sup>a</sup> $p < 0.05$ Vs normal group. <sup>b</sup> $p < 0.05$ Vs CSE exposure group. Abbreviation: CSE, cigarette smoke extract; Cur, curcumin; ML, montelukast ; AChE, acetylcholine esterase; GSH, reduced glutathione; and LPO, lipid peroxidation.

## Discussion

The present study results are revealed that pre-exposure of curcumin (10 and 20 mg/kg; 10 min/day for 7 consecutive days) shown significant (*p* < 0.05) ameliorate the effect against CSE exposure challenge induced cognitive impairment. It is indicated in various neurocognitive behavioral changes such as decreasing TSGC and raising NERC values in color recognition test (Fig. 1a and 1b); decreasing % ETC and TSTC values in partition preference test (fig 2a and 2b); decreasing TSUS and increasing the TSLS values in horizontal compartment test (Fig. 3a and 3b); and increasing TL and decreasing % TCP values in T-Maze tests (Fig. 4a and 4b). Further, the pre-treatment of curcumin also produce the potential attenuation of CSE induced biomarkers changes in the zebrafish brain. These results are statistically comparable with reference control *i.e.,* montelukast exposed group readings.

CSE consists of numerous toxic ingredients and it alters the physiological process of the body system. The primary events of CS cause the potent activation of free radicals and immune cells lead to initiates the oxidative stress, mitochondrial dysfunction and imbalance cellular oxidant and anti-oxidant balance(Tweed et al., 2012; Yun et al., 2018). The endogenous anti-oxidant, *i.e.,* reduced glutathione undergoes the lack the synthesis and rapid degradation process (Ellman, 1959). Similar results are observed in the present study, *i.e.,* decrease the GSH levels. Further, the free radical accumulation and/or generation are responsible to modulate the cell membrane integrity via activation of lipid peroxidation process(Petersen, 2017). Experimentally, it can be detected with thiobarbituric acid. Because the intermediate products of lipid peroxidation *i.e.,* malondialdehyde (MDA) readily react with thiobarbituric acid and generates pink colour chromogen (Ohkawa et al., 1979). Lipid peroxidation products detected by spectrophotometrically and results showed CSE enhance the lipid peroxidation (Chan et al., 2016). Later it also alters the mitochondrial membrane peroxidation and produces the opening of mitochondrial permeability transition pore (MPTP) viz apoptosis (Szczesny et al., 2018). In nuclear levels, it alters the nucleic acids mutation and producescancer, tumour and other ailments (Lee et al., 2018).

CSE contents posses the higher affinity to various tissue proteins such as lung, kidney, liver, heart including neuron system(Lee et al., 2018). In the brain, it alters the neuroendocrine and neurotransmitter functions lead to changes the neurobehavioural pattern (Tweed et al., 2012). In addition, the chronic exposure of CSE is known to cause the neurocognitive dysfunction via alteration of cholinergic neurotransmitter, *i.e.,* acetylcholine (Hall et al., 2016). Acetylcholine degrades by two enzymes *i.e.,* acetylcholinesterase and butrylcholinesterase leads to reduce the availability of acetylcholine content and enhances the cognitive failure (Vani et al., 2015). *In vitro* acetylcholinesterase activity estimated by using acetylthiocholine (as a substrate) and Ellman reagents (Ellman et al., 1961). The results revealthat CSE challenge enhances the acetylcholinesterase activity in zebrafish brain (Muthuraman A and Rishitha N, 2018; Rishitha and Muthuraman, 2018). These results produce the evidence of CSE challenges induced neurocognitive impairment and their modulatory action of cholinergic neurotransmitter actions. In addition, montelukast is also produced the similar results and it is established molecule in the reduction of inflammation as well as neuroprotective agents (Grinde and Engdahl, 2017; Zhang et al., 2016). Data in hand, our previous report and other research laboratory reports are collective support the curcumin possess the potential ameliorative effects in cognitive function against CSE exposure challenges. The neuroprotective mechanism of curcumin in CSE induced neurodegeneration and cognitive dysfunction illustrated in Fig. 6.

**Figure 6.**
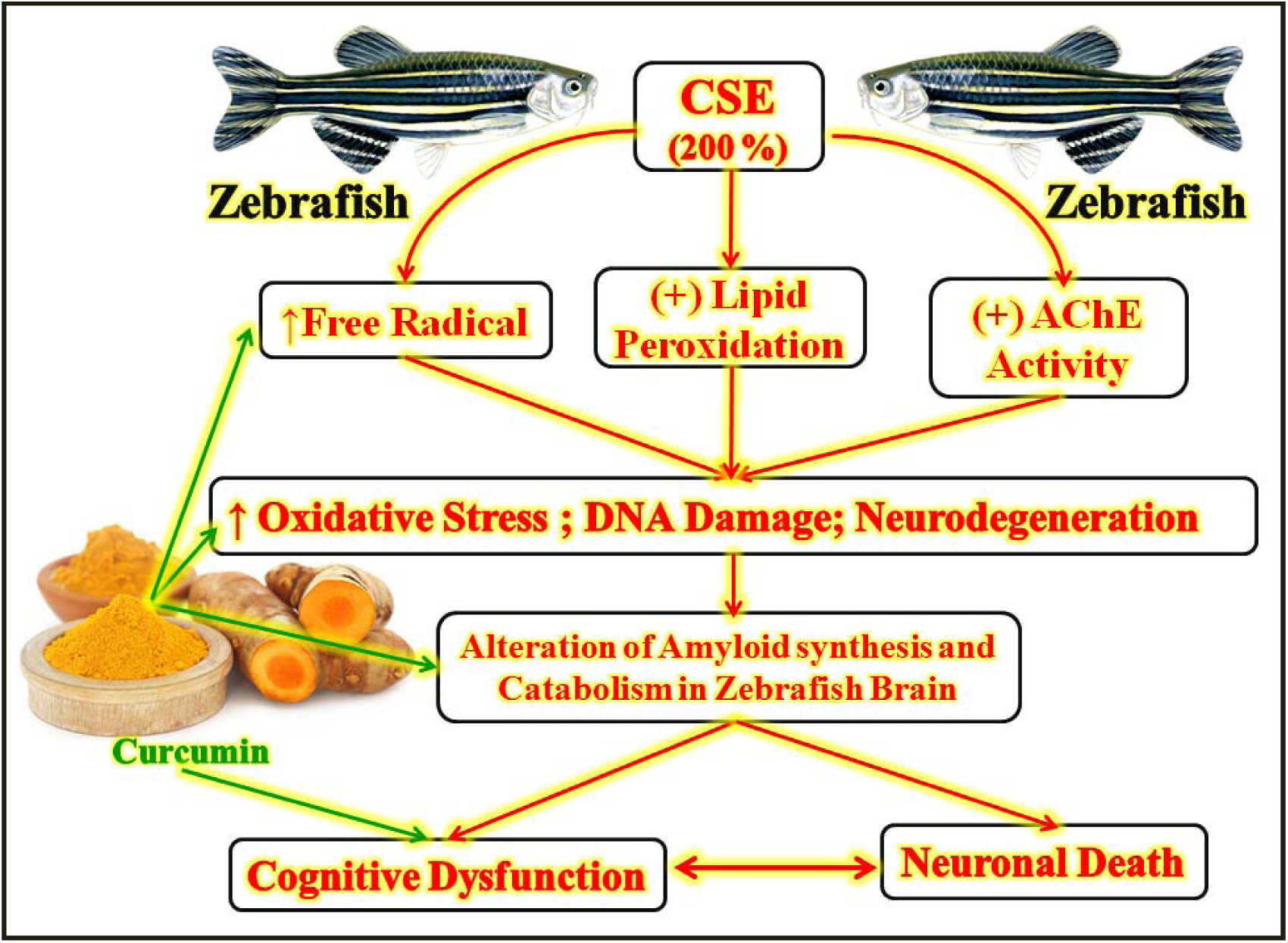
The neuroprotective mechanism of curcumin in CSE induced neurodegeneration and cognitive dysfunction. Abbreviation: CSE, cigarette smoke extract; and AChE, acetylcholinesterase.

Hence, it may conclude that curcumin may act as neuroprotective and cognitive enhancing molecule in CSE associated neuronal dysfunction and neurotoxicity due to its potential pleiotropic action, such as free radical scavenging; anti-lipid peroxidative; anti-inflammatory and neurotransmitter regulatory actions.

## ACKNOWLEDGEMENT

The authors are thankful to the Jagdguru Sri Shivarathreeshwaara University, JSS College of Pharmacy, Mysuru -570 015, Karnataka (India) for unconditional support and providing technical facilities for this research work.

## AUTHOR’S CONTRIBUTIONS

L.T., S.J., S.B.R., S.S.A., and T.G contributed in data collection and writing of the article. N.R. contributed to the design, interpretation of data and writing of the article. A.M. and N.P. contributed to the statistical analysis and critical evaluation of manuscript preparation.

